# Structural annotation of full-scan MS data: A unified solution for LC-MS and MS imaging analyses

**DOI:** 10.1101/2024.10.14.618269

**Authors:** Shipei Xing, Vincent Charron-Lamoureux, Måns Ekelöf, Yasin El Abiead, Huaxu Yu, Oliver Fiehn, Theodore Alexandrov, Pieter C Dorrestein

## Abstract

Public liquid chromatography-mass spectrometry (LC-MS) and MS imaging metabolomics data repositories contain millions of files, with over 40% of them consisting solely of MS1 (full-scan) information, creating a significant gap in data reuse potential due to limited annotation capabilities. Here, we present **ms1-id**, an open-source Python package providing a unified solution for structural annotation of full-scan MS data applicable to both LC-MS and MS imaging analyses. Our approach leverages in-source fragments to generate pseudo MS/MS spectra through correlation analysis in either chromatographic or spatial domains. We introduce precursor-tolerant reverse spectral matching that accommodates multiple ion forms simultaneously and peak intensity scaling that enables matching of low-energy in-source fragments against existing reference MS/MS libraries. Applied to inflammatory bowel disease cohorts and diverse MS imaging samples, our method uncovers metabolites previously overlooked in traditional analyses. This strategy effectively addresses a critical need in metabolomics data reuse by enabling level 2/3 structural annotation of MS1-only data, facilitating new biological insights from existing repository data that was previously only annotated at the molecular formula level.

## Introduction

The scientific community is investing substantial resources – funds, time, and effort – in collecting samples, obtaining metabolomics data, analyzing results, and making them publicly available. Just as a conservative number, we are approaching a milestone of one million public liquid chromatography-mass spectrometry (LC-MS) files and over 13,000 MS imaging files across major repositories such as Metabolomics Workbench^1^, MetaboLights^2^, GNPS/MassIVE^3^, and METASPACE^4^. The rationale behind public data deposition (even when restricted access is employed) extends beyond promoting scientific transparency and reproducibility; it also facilitates future reuse, as typically only a small fraction of the data is utilized upon initial publication.

Given that the average annotation rate in metabolomics studies ranges from 10% to 20%^5–8^, one crucial aspect of data reuse is to provide new annotations that can be re-contextualized to uncover new biological insights. In untargeted metabolomics, two primary approaches are employed: data collection with tandem MS (MS/MS) and without (MS1-only or full-scan). The latter method offers an advantage in peak shape quality due to the increased number of scans contributing to each peak, leading to enhanced absolute or relative quantification accuracy and thus more reliable statistical analyses. However, MS1-only data presents limitations in discovering molecules that were detected but not yet annotated through subsequent reanalysis. Notably, more than 40% of untargeted LC-MS metabolomics data files in public repositories consist solely of MS1 data^9^ (**Supplementary Table 1**), and almost all MS imaging metabolomics data are MS1 scans. This situation creates a significant gap in data reuse and reinvestigation for full-scan data in untargeted metabolomics.

Traditionally, MS1 data interpretation relies on accurate mass measurements and isotopic patterns, which can suggest possible molecular formulas^4,10,11^ but often falls short of providing structural information. However, it is also generally understood that many ions may undergo in-source fragmentation or exhibit post-source decay^12–17^, generating fragment pieces that also appear in MS1 data. As these processes involve thermal activation, the resulting in-source fragment ions exhibit fragmentation patterns very similar to those observed in collision-induced dissociation (CID) MS/MS spectra. This opens the possibility of leveraging such in/post-source fragments as a handle to create pseudo MS/MS spectra, also referred to as composite spectra^18^, that can be leveraged for MS/MS reference library-based annotation in metabolomics and exposomics studies^16–24^.

Strategies, such as IDSL.CSA^18^, have demonstrated the proof-of-principle of matching pseudo MS/MS spectra, obtained by aggregating ion forms across entire datasets, to reference MS/MS libraries using scoring methods like cosine or entropy similarity^25^. This works particularly well for GC-MS^26^ due to the consistent use of a fixed energy (70 eV) for both data acquisition and reference spectra, and the absence of many co-eluting ion forms such as different adducts or multi-meric species. However, in LC-MS, different ion forms, such as adducts and multimers, often dominate pseudo MS/MS spectra, which may prevent matching to reference MS/MS libraries that do not account for these ion forms. Furthermore, as we show below, the fragment ions we detected tended to match reference MS/MS spectra generated with lower-energy fragmentation, while most reference spectra available in the public domain are collected under medium to high collision energies. In these lower-energy spectra, low-*m/z* ions often appear at very low intensities or may be absent entirely, resulting in missed matches. Other experimental strategies aimed to overcome this limitation–for example, eISA^27^ and EISA-EXPOSOME^19^ have been developed to incorporate in-source fragments in metabolite annotation. Therefore, in their current implementations, these methods require full-scan data under enhanced in-source fragmentation energies (e.g., 40 eV) to obtain adequate spectral matches, and they work in a targeted fashion. Consequently, these methodologies cannot be used for reanalyzing hundreds of thousands of public untargeted MS1 data files acquired without extra in-source fragmentation (referred to as “in-source collision-induced dissociation” or isCID in MS instrument settings) experimental designs.

In the realm of MS imaging, metabolite candidate annotations are obtained by annotating molecular formulas and cross-referencing these formulas in common metabolite/lipid chemical databases such as HMDB^28^, LipidMaps^29^ and ChEBI^30^. This approach corresponds to level 4 identification confidence (unequivocal molecular formula) according to the Metabolomics Standards Initiative^31^. Despite the rapid growth of MS/MS spectral libraries^8^, metabolite annotation of MS imaging data has not yet fully benefited from this expanding community resource.

In this work, we present an approach for comprehensive MS1 data annotation applicable to both LC-MS and MS imaging analyses. Our method integrates correlation-based feature clustering with reverse spectral matching strategies to leverage in-source fragments for structural annotation. We introduce three key innovations: (1) a unified computational framework that treats both chromatographic and spatial domains equivalently for pseudo MS/MS generation; (2) precursor-tolerant reverse matching that simultaneously accommodates multiple ion forms during spectral matching; and (3) a mass-dependent peak intensity scaling approach that enables matching low-energy in-source fragments against existing reference MS/MS libraries collected at medium to high collision energies. This approach not only provides structural information for MS1-only LC-MS data but also represents the first implementation of MS/MS library-based structure annotation for MS imaging data. Our open-source Python package, **ms1-id**, facilitates full-scan MS data annotation across analytical platforms, unlocking new biological insights from the vast repositories of existing data that previously could only be annotated at the molecular formula level.

## Results

Our approach integrates two steps (although how these steps are implemented is critical): (1) clustering ions or metabolic features through correlation analysis of extracted ion chromatograms (XICs) or ion images in the retention time domain or spatial manner (**Fig. 1a**); and (2) employing a precursor-tolerant (open search) but using reverse spectral matching approach to compare deconvolved MS1 spectra, or pseudo MS/MS spectra, against peak scale-adjusted reference MS/MS libraries for structure candidate identification. Unlike traditional forward spectral matching which utilizes all the peaks in both query and reference spectra for scoring, reverse matching is a unidirectional spectral comparison which ignores unaligned peaks in the query spectrum^32^, tolerating contaminant peaks sourced from co-eluting metabolic features or signal artifacts. In the following sections, we elaborate on the spectral matching design and its underlying rationales, demonstrating how this approach enhances annotation capabilities for MS1 data in both LC-MS and MS imaging experiments.

**Fig. 1.**
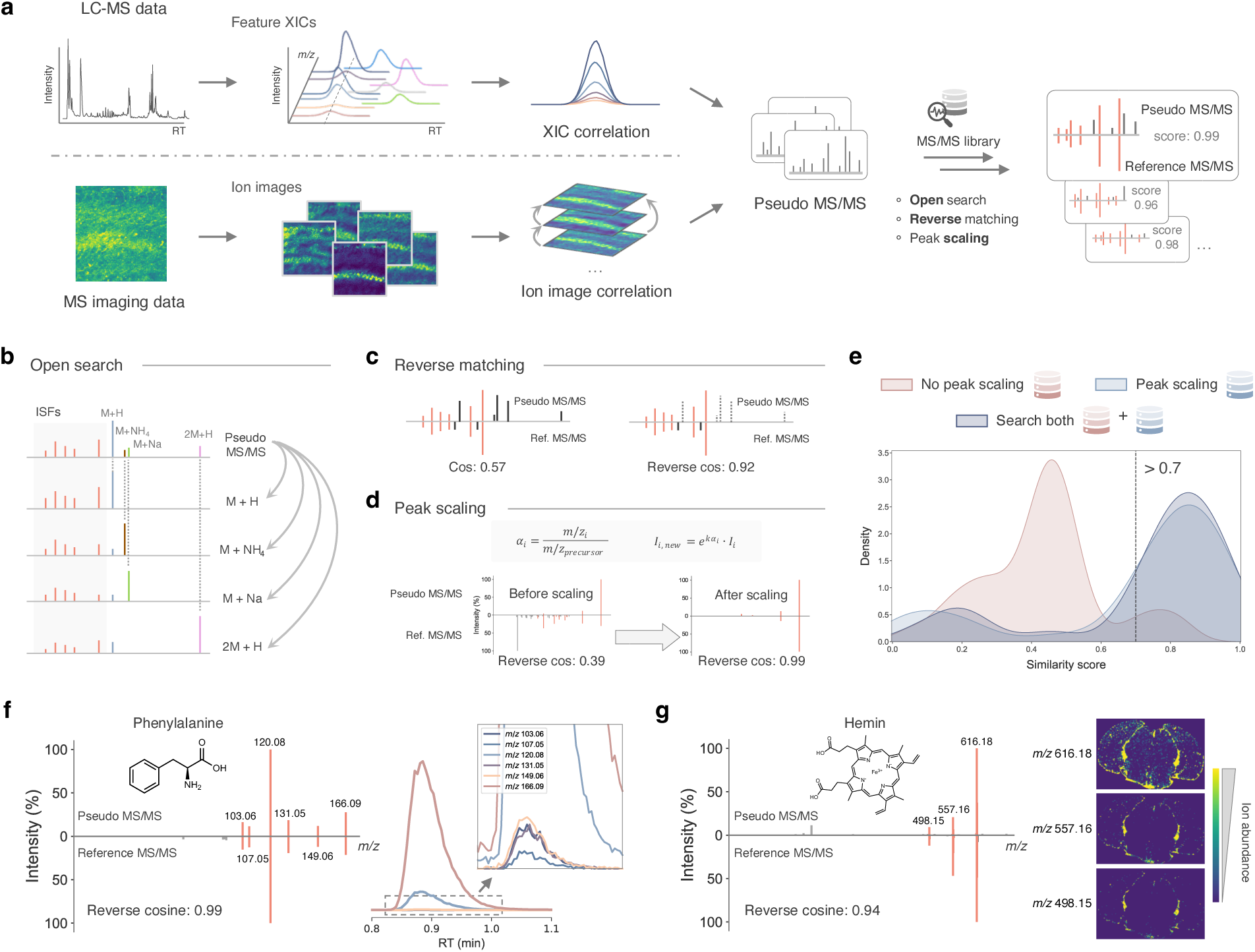
Structure annotation of full-scan MS data. **a**, A unified solution to annotate MS1 data from LC-MS or MS imaging experiments. Pseudo MS/MS spectra are generated through correlation analyses in time or spatial domain. The precursor ion-tolerant (open search) reverse spectral searching allows structure annotation of pseudo MS/MS leveraging existing reference MS/MS libraries. **b**, Open search allows matching against reference spectra of multiple adduct forms. ISFs, in-source fragments. **c**, Reverse spectral search discards unmatched peaks in the query spectrum and thus improves spectral search. **d**, Peak intensity scaling helps to match pseudo MS/MS spectra against reference MS/MS which are collected under medium to high collision energies. **e**, Similarity score distribution of searching pseudo MS/MS against libraries with or without peak scaling, or both. Reverse cosine scores of ground truths using chemical standards are used. **f**, An example of structure annotation from LC-MS data (NIST human feces) and XIC correlations. Peak intensities are square rooted. **g**, An example of structure annotation from MS imaging data (mouse brain) and extracted ion images. Ion images were created using 5 ppm mass tolerance. Peak intensities are square rooted.

Each molecule detected by mass spectrometry is represented by multiple molecular ions of various adduct forms (e.g., [M+NH_4_]^+^, [M+Na]^+^) that co-elute during chromatography^33–35^ or, in the case of MS imaging, share spatial correlations with each other, in addition to in-source fragments. Consequently, intact molecular ions of different adducts, along with their fragments, appear in the same reconstructed pseudo MS/MS spectrum (**Fig. 1b**). Unlike a typical MS/MS spectrum collected in data-dependent acquisition (DDA) mode, it is not known which ion–if any– represents the precursor ion in a pseudo MS/MS. To address this, we implemented an open search approach–it does not assume a single, predefined precursor ion for each spectrum, but instead considers every ion as a potential precursor ion simultaneously. It employs an unlimited mass tolerance window to accommodate potential mass shifts due to different adducts or multimers, enabling the recognition of various precursor types within the same spectrum.

However, there are challenges in matching a pseudo MS/MS spectrum against the MS/MS spectral library, making it often not possible to provide direct matches. Pseudo MS/MS spectra contain not only fragment ions but also molecular ions of other adduct types and unavoidably mis-clustered ions that are co-eluting. These additional ions are undesirable during spectral matching as they significantly diminish the search scores as the reference libraries do not contain all of the different ion forms. We therefore employed the reverse spectral search^36^, an algorithm first introduced by Abramson in 1975 and initially designed to overcome interference in complex GC-MS data. The key innovation of reverse search is its ability to selectively focus on diagnostic peaks while ignoring interference, making it particularly well-suited for complex spectra containing multiple ion species. Applying this concept to our pseudo MS/MS data, reference spectra serve as templates, and unmatched peaks in the pseudo MS/MS are discarded when calculating matching scores (**Fig. 1c**). This approach is particularly crucial for MS imaging data, where ions with similar biological functions tend to have similar spatial patterns and thus high correlations, resulting in more ions that should not be compared when trying to annotate (e.g., lipid molecules can exhibit similar spatial distributions on cellular membranes).

Furthermore, as pseudo MS/MS spectra are obtained with minimal energy input (only enough for ion transfer and/or trapping), the fragment ion intensities tend to more closely align with low-energy CID spectra. Currently, most reference MS/MS spectra in libraries are collected under medium to high collision energies (**Supplementary Fig. 1**). Analysis on the MoNA library revealed that <1% of the reference spectra were acquired under collision energies of ≤5 eV. Therefore, we developed a peak intensity scaling approach to better align them (**Fig. 1d**). Using chemical standard pools of bile acids and drugs, in total containing 14 known molecules, for which full-scan MS data were collected under in-source fragmentation energies of 0 eV, 10 eV and 20 eV, we demonstrated that this peak scaling approach provided more matching scores of >0.7 for ground truths compared to not applying peak scaling (**Fig. 1e**). Combining search results from both original and peak-scaled reference libraries maximized the number of matches with reverse cosine scores exceeding 0.7. However, by analyzing low-energy reference MS/MS spectra from the NIST20 library against the GNPS library (**Supplementary Fig. 2**), we observed that the peak scaling approach can increase the false discovery rate (FDR) during spectral matching. We anticipate future algorithmic refinements will enable more stringent quality control while maintaining high annotation rates.

To further validate our approach, we collected LC-MS data from NIST reference human fecal samples in both DDA and full-scan modes. Full-scan data were acquired under 0 eV, 10 eV, and 20 eV in-source fragmentation energies. We were able to obtain spectral library matches for 567, 306, 484, 511 and 604 metabolic features in MS/MS (DDA, 42 eV), MS1 (DDA), MS1 (0 eV), MS1 (10 eV), and MS1 (20 eV) modes, respectively (**Fig. 2a**). Unexpectedly, MS1 annotation revealed a unique chemical space, with the majority of annotated features in MS1 data being distinct from MS/MS annotations. More than 79% of the features annotated via pseudo MS/MS lacked corresponding MS/MS spectra in DDA experiments. While DDA typically acquires MS/MS spectra for the more abundant features, this approach captures more low-intensity features when they produce sufficient in-source fragments (**Fig. 2b**). When examining the same metabolic features collected in DDA, structure similarity analyses between MS1 annotations and MS/MS annotations showed that they generated similar chemical candidates (**Fig. 2c**), where a higher in-source fragmentation energy led to more similar or identical structure matches with MS/MS annotations. We then investigated the compound classes^37^ of annotated compounds under different acquisition conditions (**Fig. 2d**). While MS1 data generally annotated more molecules than MS/MS across most compound classes, organic acids & derivatives, and lipids & lipid-like molecules were not as well recognized in MS1 annotation compared to MS/MS, and this suggests that certain classes of compounds will be easier to annotate via the pseudo MS/MS strategy forwarded here. Overall, above results indicate that MS1 annotation expands the range of detectable metabolites, potentially uncovering previously overlooked compounds in untargeted metabolomics studies, including those available in public repositories.

**Fig. 2.**
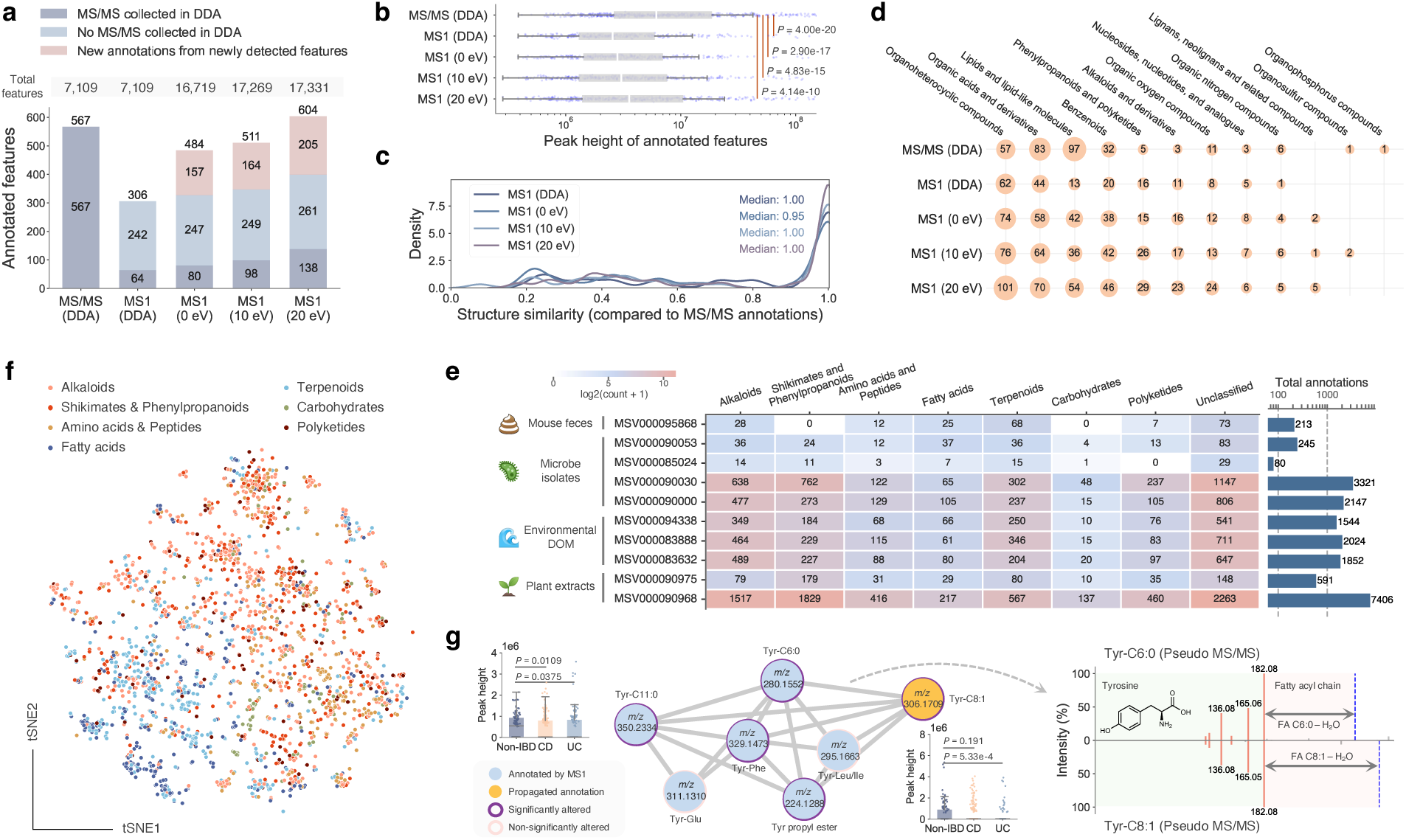
MS1 annotation of LC-MS data using MS/MS libraries. **a**, Annotations of NIST fecal samples by MS/MS (DDA) or MS1 data with different in-source fragmentation energies. **b**, MS1 annotation is able to capture low-abundant metabolic features compared to MS/MS annotations in DDA. Boxes cover the interquartile range (IQR), with medians labeled. The upper whisker represents Q3 + 1.5 × IQR; the lower whisker represents Q1 − 1.5 × IQR. *P* values were calculated using two-sided Mann-Whitney *U* tests. **c**, Structure similarity distributions between MS1 annotations and MS/MS annotations when annotating the same metabolic features. **d**, Compound class distributions of the metabolites annotated in the NIST feces dataset. **e**, Annotation summary of 10 public untargeted metabolomics datasets selected from diverse sample types. **f**, t-SNE visualization of MS1 annotations in the IBD dataset. Nodes are colored by compound pathways from NPClassifier. **g**, An example molecular network generated using pseudo MS/MS spectra in the IBD dataset (HILIC positive). Tyrosine-related compounds were annotated and linked. *P* values were calculated using two-sided Mann-Whitney *U* tests. The mirror plot shows pseudo MS/MS spectra from Tyr-C6:0 and Tyr-C8:1.

To demonstrate the robustness and broad applicability of our approach, we reanalyzed 10 public LC-MS untargeted metabolomics datasets spanning diverse sample types, including mouse feces, microbial isolates, environmental dissolved organic matter (DOM), and plant extracts (**Fig. 2e**). These datasets encompass various LC-MS platforms, instrument types, and ionization polarities. Our method successfully annotated a wide spectrum of compound classes, with particularly strong representation of alkaloids, shikimates & phenylpropanoids, and terpenoids across the different matrices.

As a further validation on clinically relevant data, we revisited a public LC-MS full-scan dataset from an inflammatory bowel disease (IBD) study^38^. This dataset comprised 546 stool samples from three diagnostic groups: non-IBD (n = 134), Crohn’s disease (CD, n = 266), and Ulcerative colitis (UC, n = 146). The analysis employed four distinct LC-MS modes: HILIC positive, HILIC negative, C8 positive, and C18 negative. We performed MS1 annotations in batches, obtaining 3010, 293, 227 and 636 unique metabolites (unique InChIKey strings) in the four modes, respectively. Altogether, we identified 3802 unique metabolites with level 2/3 confidence^31^. A t-SNE visualization (**Fig. 2f**), color-coded by compound pathways from NPClassifier^39^, revealed distinct clustering patterns among annotated compounds. Alkaloids and shikimates & phenylpropanoids form large, spread-out clusters, suggesting diverse structural variations and their prominence in the gut metabolome. Fatty acids and terpenoids form relatively distinct clusters, indicating unique intensity profiles for lipid-based metabolites across the IBD sample set. We constructed a molecular network using the pseudo MS/MS spectra from the HILIC positive mode. A subnetwork for tyrosine-related compounds was highlighted in **Fig. 2g**, including N-acyl amides that showed alterations in non-IBD vs CD or non-IBD vs UC comparisons. The annotation of Tyr-C8:1 could be propagated through modified cosine-based MS/MS similarity and delta masses with neighbor nodes, and the mirror plot clearly shows the fragmentation pattern of tyrosine as well as the neutral loss of the fatty acyl chain. These findings align with previous IBD studies^40–42^, which identified alterations in lipid metabolism and N-acyl amide profiles as key factors in IBD pathogenesis, and highlight that our reanalysis approach can assist in uncovering clinically relevant metabolic signatures the original depositors did not describe.

To validate our annotation approach on MS imaging data, we analyzed data that had seven spotted chemical standards (ATP, ADP, adenosine, guanine, glutamic acid, glutamine, and glutathione) on mouse liver tissue (**Fig. 3a**) and collected imaging data in negative ion mode. We then assessed whether these compounds could be retrieved using our pseudo MS/MS approach. Among the seven spotted standards, five were successfully annotated. Guanine (*m/z* 150.0421) and glutamic acid (*m/z* 146.0459) could not be confidently annotated as they did not generate sufficient in-source fragment ions within our collected *m/z* range (*m/z* > 100). Using ATP as a representative example, we observed multiple co-localized in-source fragments (**Fig. 3b-c**), including sequential losses of phosphate groups: [ATP-HPO_3_-H]^-^, [ATP-2HPO_3_-H]^-^, [ATP-2HPO_3_-H_2_O-H]^-^, and the phosphate chain fragment [H_3_P_3_O_10_-H]^-^. The spatial correlation between these in-source fragments and the parent ATP molecule not only confirmed their structural relationship but also demonstrated how our method leverages these fragment patterns for confident annotation.

**Fig. 3.**
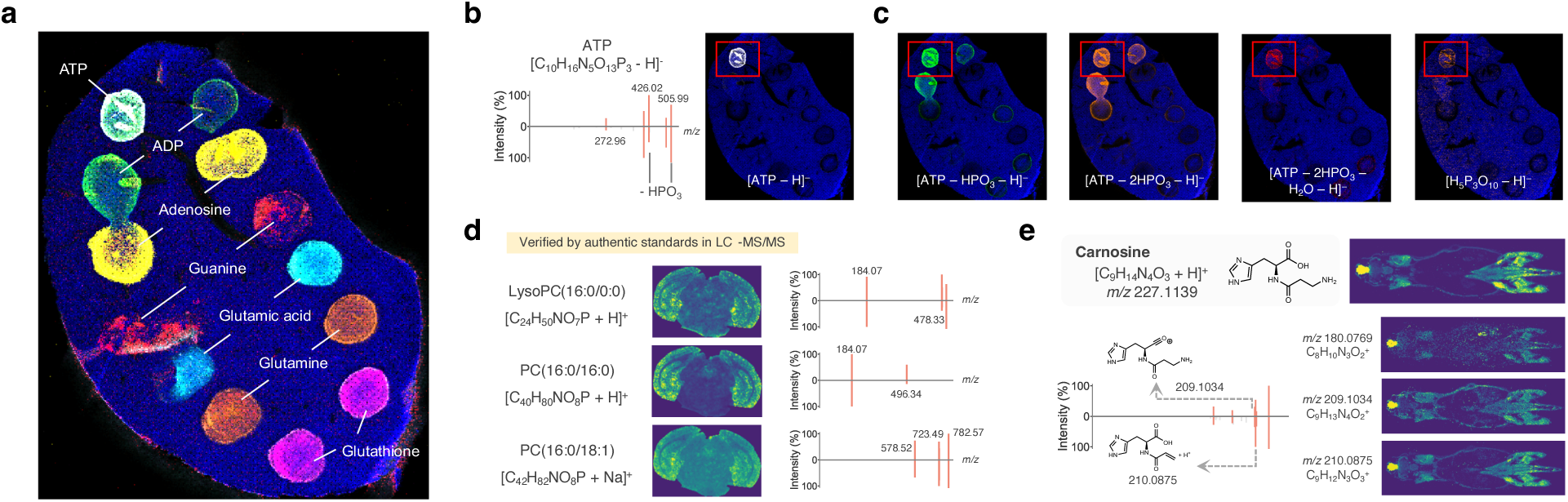
Annotation of MS imaging data using MS/MS libraries. **a**, Overlay of ion images from seven chemical standards (ATP, ADP, adenosine, guanine, glutamic acid, glutamine, and glutathione) spotted on the mouse liver tissue. **b**, Mirror plot showing ATP annotation with its corresponding ion image. **c**, Ion images of in-source fragments co-localized with ATP, demonstrating the spatial correlation of parent molecule and fragments. **d**, Phospholipid annotations in mouse brain imaging data: LysoPC(16:0/0:0), PC(16:0/16:0) and PC(16:0/18:1), all verified by authentic standards in LC-MS/MS data. Mirror plots show pseudo MS/MS spectra with characteristic fragment ions at *m/z* 184.07 (phosphocholine head group). **e**, Carnosine annotation in mouse body imaging data showing localization in brain and muscle tissues with corresponding fragment ion images. All ion images were created using 5 ppm mass tolerance. Peak intensities are square rooted in mirror plots, with matched peaks shown.

To further demonstrate the efficacy of our strategy on MS imaging data, we applied it to multiple datasets including mouse brain^4^, mouse body, mouse kidney, human kidney and plant root. In the mouse brain sample, we annotated hemin and phosphatidylcholine (PC) lipids of varying chain lengths. **Fig. 1g** displays the ion images of the hemin cation and its in-source fragments, and the visual inspection clearly revealed that the ion image of the hemin cation exhibits spatial patterns highly similar to those of its in-source fragments, with the expected lower abundance of in-source fragments. **Fig. 2g** illustrates the annotations of LysoPC(16:0/0:0), PC(16:0/16:0) and PC(16:0/18:1), which were all verified by authentic standards in LC-MS/MS data^4^. In addition, our method successfully handles different adduct forms of the same molecule, preserving their distinct biological distributions and fragmentation patterns. **Supplementary Fig. 3** demonstrates this capability with PC(16:0/18:1) annotated in both [M+H]^+^ and [M+Na]^+^ forms (reverse cosine scores 0.85 and 0.77, respectively). The ion images clearly show distinct spatial distributions: the [M+H]^+^ form predominates in the cortex and hippocampus regions, while the [M+Na]^+^ form shows higher intensity in midbrain regions and white matter areas. In the mouse body dataset, we annotated carnosine, which was localized to the brain and muscle tissues (**Fig. 2h**), aligning with its dual role as a neuroprotector and a muscle performance enhancer^43^. In the brain, carnosine’s presence suggests its involvement in neurotransmitter regulation and synaptic plasticity, processes crucial for learning and memory^44^. In muscle tissue, it functions as an intracellular buffer, regulating pH levels during physical activity, and exhibits antioxidant properties^45^ that may aid in recovery from exercise-induced stress. The significant abundance of carnosine in these tissues underscores its importance in both neurocognitive function and physical performance. In the kidney samples, we identified several energy metabolism-related compounds including adenosine derivatives (NAD+, adenosine diphosphate) and flavin mononucleotide. In the plant root samples, we annotated key primary metabolites including phosphorylated sugars (glucose-1-phosphate, fructose-6-phosphate) and organic acids (citric acid, isocitric acid, malic acid), which are central components of plant metabolism.

Extending our analysis to a single cell dataset of human cell lines, we examined an MS imaging dataset from differentiated human hepatocytes^46^ revealed various lipid classes including phosphatidylcholines, diacylglycerols, and triacylglycerols. As an illustrative example, **Supplementary Fig. 4** showcases the annotation of a diacylglycerol species. This result highlights the ability of this approach to annotate complex lipids that are interpreted at the single cell level. In another single cell dataset (HeLa and mouse fibroblast cells)^46^, we primarily observed phospholipid species including phosphatidylethanolamines (PEs) and phosphatidylinositols (PIs), reflecting the cellular membrane composition of cultured cells. Additionally, we applied our approach on MALDI-TOF MS imaging datasets, which are widespread but often present challenges due to lower resolution and mass shift issues. Using mouse brain and human liver MALDI-TOF datasets, we successfully annotated small metabolites (e.g., levodopa, CoQ10) and various lipid species, extending the utility of our method to legacy imaging data. These findings collectively demonstrate the versatility of our approach across different types of data, from tissue-level imaging to single-cell analysis, and across various MS platforms and sample types. By enabling confident annotation of molecular species in various biological contexts, our method promises to enhance our understanding of spatial metabolomics and lipidomics in health and disease.

## Discussion

Other than tools such as IDSL.CSA^18^, eISA^27^, EISA-EXPOSOME^19^ that we mentioned earlier, there are several other approaches leveraging in-source fragmentation for metabolite annotation that should be discussed here. HERMES^47^ utilizes public low-energy MS/MS data to annotate in-source fragments, filtering ionic formula features that correspond to artifacts rather than endogenous metabolites. This approach is primarily designed to optimize targeted MS/MS acquisition by creating a high-quality inclusion list after removing in-source fragments. In contrast, our approach leverages in-source fragmentation as diagnostic information, generating pseudo MS/MS spectra from MS1 data to enable direct annotation. This fundamental difference makes our method particularly valuable for reanalyzing existing MS1-only datasets where further data acquisition is not possible.

For MS imaging data specifically, METASPACE^4^ uses a target-decoy FDR estimation approach for molecular formula determination (MSI^31^ level 4 confidence), while our approach aims to provide structural information (MSI level 2/3) by leveraging spectral matching of in-source fragments. When comparing our results from the mouse brain dataset with METASPACE annotations, we found that the approaches are highly complementary. While METASPACE excels at formula-level annotation, our method can distinguish between isomeric structures that share the same formula and can identify in-source fragments that might otherwise be misannotated as intact metabolites. This complementarity is particularly valuable for phospholipids like PCs and PEs, where in-source fragmentation is common and can lead to incorrect annotations in formula-only approaches.

Moreover, rMSIfragment^48^ is a specialized solution for annotating in-source fragments in MALDI-MSI lipidomics. This exploits known adducts and fragmentation pathways to confidently annotate lipids in MS imaging data. As a complementary approach, **ms1-id** provides broader coverage across metabolite classes and is not limited to lipids or specific analytical techniques. The integration of rMSIfragment’s lipid-specific knowledge with our broader metabolite coverage could further improve annotation confidence in lipidomics studies.

Despite its capacity to annotate MS1 data from both LC-MS and MS imaging experiments, there are a number of important limitations one has to consider when applying our approach. This strategy will not be able to distinguish most isomers, particularly in complex metabolite mixtures with inadequate chromatographic separation. These structurally similar compounds often co-elute and produce similar fragments, impeding the creation of clean pseudo MS/MS spectra and their subsequent distinction. Another limitation relates to the handling of different adduct forms. When compounds predominantly form adducts like [M+NH_4_]^+^ during ionization, their in-source fragmentation patterns may not match available reference spectra, which are primarily collected for protonated and deprotonated forms. This particularly impacts certain compound classes that preferentially form specific adducts, such as lipids and carbohydrates. While molecular networking or modified cosine similarity^49^ can help by capturing related spectra when adduct shifts are reflected in fragment ions, this remains a fundamental challenge for library-based annotation approaches.

Pseudo MS/MS spectra, which are reconstructed from MS1 data, tend to match most closely with low-energy reference spectra. However, as low-energy spectra are underrepresented in public spectral libraries, we proposed a peak scaling approach to fully leverage existing spectral libraries. While current peak scaling parameters are optimized to maximize annotation coverage, this optimization comes with an increased rate of incorrect matches compared to non-scaled approaches (**Supplementary Fig. 2**). We expect future optimization of scaling can further improve the annotation confidence. Another consideration is the potential for ion contamination or incorrect ion clustering when generating pseudo MS/MS spectra, especially in MS imaging data lacking chromatographic separation. Such limitations elevate the risk of incorrect matches to reference MS/MS spectra. As we show, certain compound classes are underrepresented (e.g., organic acids and lipids) in MS1 annotation. This underrepresentation arises from insufficient generation of in-source fragments due to the comparatively low energy imparted on the ions in MS1-only scans, and/or the failure of data collection of low *m/z* range (50 to 200). For MS imaging data, we recommend including the low *m/z* range to fully inspect the in-source fragments, which can be better utilized for MS1 annotation. These constraints highlight avenues for future research, including the advancement of more precise MS1 data deconvolution techniques, incorporation of additional orthogonal data for isomer differentiation, and refinement of spectral search algorithms specifically tailored for MS1 data annotation.

While our package, **ms1-id**, currently implements MassCube^50^ for metabolic feature extraction of LC-MS data in the workflow, **ms1-id** is designed to be compatible with pseudo MS/MS spectra generated by various established tools such as CAMERA^33^, CliqueMS^51^, and xMSannotator^52^. Users can generate pseudo MS/MS spectra using their preferred tool and utilize the **ms1-id** package for annotation through its command-line interface. For computational efficiency, **ms1-id** processes LC-MS data files in parallel using multiprocessing, while MS imaging data processing follows a sequential workflow optimized for memory management when handling large spatial datasets. As practical benchmarks, using 6 cores on macOS (M2 Max), processing 6 mzML files (262 MB) takes 29 minutes for feature extraction, pseudo MS/MS generation, and annotation. For MS imaging data, processing one mouse brain dataset (imzML: 29 MB; ibd: 56 MB) takes 4 minutes using 12 cores on the same computer. Memory requirements scale with dataset size, with typical analyses requiring 4-8 GB RAM.

Our MS1 annotation approach unveils exciting new prospects for untargeted metabolomics data reuse and reanalysis. A key opportunity lies in developing an MS1-based MASST^53,54^ (Mass Spectrometry Search Tool) to perform reverse metabolomics^55^ on LC-MS and MS imaging data, which allows the contextualization of molecules (known or unknown) driven by metadata integration^9^ including body distributions, producing organisms, health conditions and interventions. While the current MASST enables searching MS/MS spectra against public data repositories using forward (modified) cosine to retrieve valuable metadata for new biology discovery, MASST could now potentially be extended to the MS1 level. As a proof-of-principle, we queried the pseudo MS/MS spectrum of phenylalanine-C3:0 from the NIST feces sample, which was more abundant in the omnivore group than the vegan group, against the pseudo MS/MS spectra pool from the IBD dataset. This search returned an MS/MS match with cosine score of 0.90 (**Supplementary Fig. 5**). The matched pseudo MS/MS was also annotated as phenylalanine-C3:0 in the IBD dataset, showing statistical significance in both non-IBD vs CD and non-IBD vs UC comparisons with higher abundance in the non-IBD group. This indicates the feasibility of MS1-based MASST across all four major repositories. Our MS1 annotation approach’s ability to identify low-abundance features suggests the possibility of achieving broader metabolome coverage through MS1-based molecular networking^3^. This approach could catalyze the propagation of annotations through spectral similarity analysis, revealing previously unidentified metabolites and facilitating the creation of pseudo MS/MS-based suspect libraries^56^ for future data reuse and reanalysis. With over 15,000 untargeted metabolomics datasets (~one million data files) currently available in public repositories, this represents an untapped resource for exploring the dark metabolome^5^–including those elusive metabolites that have thus far escaped identification. As we refine and extend our MS1 annotation techniques, we anticipate an extensive deepening of our understanding of complex metabolic processes and their roles in diverse biological systems and disease states.

## Methods

### ms1-id package

The **ms1-id** Python package is available through the Python Package Index (PyPI). Source codes are available on https://github.com/Philipbear/ms1_id. The package supports multiple input formats for MS1 data annotation:

1. Pseudo MS/MS spectra in MGF format
2. LC-MS data in mzML or mzXML format
3. MS imaging data in imzML and ibd format

Several critical parameters can substantially influence processing efficiency and annotation quality. For LC-MS data, the ‘mass_detect_int_tol’ parameter defaults to 20,000 for Orbitrap instruments and 500 for TOF instruments. Users should optimize this parameter to achieve appropriate mass detection intensity tolerance and prevent excessive computation time. Similarly, for MS imaging data, the signal-to-noise factor (’sn_factor’) defaults to 3.0, which is suitable for most applications, though adjustment may be necessary for optimal results.

### Pseudo MS/MS spectra generation

For LC-MS data, metabolic features are extracted using the MassCube^50^ backend (ver 1.0.11), which is a Python-based framework for untargeted metabolomics. For each pair of metabolic features within the same retention time window (e.g., ±1.5 s), peak-peak correlation is calculated using their chromatographic profiles. To perform the correlation analysis between two ions, they must share at least 4 consecutive MS1 scans in their chromatographic profiles.

Pseudo MS/MS spectra are then generated as follows: For each metabolic feature (target feature), all other features with correlations exceeding a predefined threshold (e.g., Pearson correlation coefficient ≥ 0.80) are collected. These correlated features are compiled into a pseudo MS/MS spectrum for the target feature. Peak heights of the correlated features in the original MS1 data are used as their respective intensities in the pseudo MS/MS spectrum. Peaks that are determined as isotope peaks by MassCube are excluded from pseudo MS/MS generation.

For MS imaging data, the process is adapted to account for spatial information. Each MS scan undergoes noise reduction using a moving average algorithm. Within a moving window of 100 Da, the baseline is determined as 5 times the mean intensity of the lowest 5% ions in the window, effectively removing background noise. Data centroiding is performed if necessary to reduce data complexity. Ion images are extracted using a predefined mass tolerance (e.g., 10 ppm), and then spatially correlated. A minimum of 50 shared pixels with non-zero intensities between two ion images is required to ensure meaningful correlations and mitigate the impact of sparse data. Pseudo MS/MS spectra are generated by applying a predefined spatial correlation cutoff (e.g., 0.85). Both Numba^57^ acceleration and parallel processing are employed for computation efficiency enhancement.

### Reverse spectral search

Reverse spectral search is an asymmetric matching process, where one spectrum is treated as template (***T***) and the other as query (***Q***). All the peaks in the template spectrum and aligned peaks in the query spectrum are involved in matching score calculation, shown as follows.

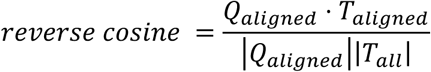

Considering that pseudo MS/MS spectra are generated from low-energy fragmentation scan modes, and that most public reference MS/MS are acquired under medium to high collision energies, we propose a mass-dependent approach to scale peak intensities for reference MS/MS spectra. This method aims to simulate the pattern observed in low-energy MS/MS, where fragment ions with *m/z* values closer to the precursor *m/z* exhibit higher abundance, while those further from the precursor *m/z* show lower abundance. For a reference MS/MS, we have

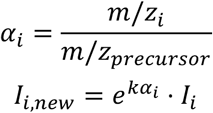

where *α_i_* is the *m/z* ratio of fragment ***i*** to the precursor; *I_i_* is the original intensity; *I_i,new_* is the scaled peak intensity; *k* is the scaling factor where we set it as 10 throughout this paper. Square root transformation is then applied on both reference and pseudo MS/MS spectra. Each pseudo MS/MS is searched against the reference MS/MS library in a precursor-tolerant manner, where we ask that the precursor ions of matching hits should be in the *m/z* values of the query pseudo MS/MS spectrum. For each pseudo MS/MS, we reserve the top 1 hit for each unique precursor *m/z* value among all annotations. Each annotation is then linked to a single metabolic feature in the metabolic feature table using the retention time of the pseudo MS/MS and the precursor mass of the annotation.

To speed up the process of library search, we modified the flash entropy framework^58^ specifically for reverse cosine search, which outputs reverse cosine score, matched peak number and spectral usage (sum intensities of matched peaks over total intensities in the query spectrum). The following cutoffs were used for LC-MS MS1 data annotation: minimum score, 0.7; minimum matched peaks, 3; minimum spectral usage, 0.20. For MS imaging data, we used: minimum score, 0.7; minimum matched peaks, 3; minimum spectral usage, 0.05.

The reference MS/MS library needs to be preprocessed and indexed before use. We provided the indexed version of GNPS MS/MS library (downloaded on July 17, 2024) as well as the code to index an MS/MS library on GitHub (https://github.com/Philipbear/ms1_id).

### Preparation of chemical standards

For the bile acid pool, a stock solution of 10 mM of glycocholic acid (GCA), glycochenodeoxycholic acid (GCDCA), taurodeoxycholic acid (TDCA), taurocholic acid (TCA), taurolithocholic acid (TLCA), and tauro-3alphahydroxy-12ketocholanoic acid was prepared. All bile acids were diluted to 10 µM into a single 2 mL LC-MS glass vial (Thermo Fisher) to create a pooled sample. For the drug pool, a stock solution of 10 mM of sertraline, venlafaxine, ritonavir, darunavir, losartan, quetiapine, sulfasalazine, and abacavir was prepared. All drugs were diluted to 10 µM into a single 2 mL LC-MS glass vial (Thermo Fisher) to create a pooled sample.

### Preparation of NIST reference materials

NIST fecal reference materials (two vegan tubes and two omnivore tubes) were subjected to a biphasic extraction^59^ to remove lipids and retain the metabolite fraction. One mL of NIST fecal material was transferred to a 2 mL Eppendorf tube and dried overnight in a CentriVap. Dry materials were resuspended with 325 µL of cold MeOH (LC-MS grade, Thermo Fisher), vortex for 10 s, and sonicated for 5 min before adding 1083 µL of cold MTBE. Samples were vortexed for 10 s and sonicated for 2 min followed by 1 h incubation at 4 °C. To induce phase separation, 271 µL H_2_O (LC-MS grade, Thermo Fisher) was added to the samples and centrifuge at 10,000 x g for 10 min. The upper phase was removed and 1084 µL of MeOH was added, followed by an overnight incubation at −20 °C. Samples were centrifuged at 15,000 x g for 10 min. An equal amount (50 µL) of the fecal NIST materials were combined to generate a pooled NIST reference fecal sample. All samples were dried in a CentriVap and stored at −80 °C until resuspension. NIST reference fecal materials were resuspended in 200 µL of 50% MeOH/H_2_O with sulfadimethoxine as internal standard before LC-MS analysis.

### LC-MS analysis

The chromatographic separation was done on a reverse phase polar C18 (Kinetex Polar C18, 100 mm x 2.1 mm, 2,6 µm, 100 angstrom pore size with the matching guard column, Phenomenex) using a Vanquish UHPLC coupled to an Orbitrap mass spectrometer (Thermo Fisher Scientific). Five microliters of samples were injected into the mobile phase, which is composed of solvent A (H_2_O with 0.1% formic acid) and solvent B (ACN with 0.1% formic acid) with the column compartment kept at 40 °C. Samples were eluted at a flow rate of 0.5 mL/min using the following gradient: 0 min, 5% B; 1.1 min, 5% B; 7.5 min, 40% B; 8.5 min, 99% B; 9.5 min, 99% B; 10 min, 5% B; 10.5 min, 5% B; 10.75 min, 99% B; 11.25 min, 99% B; 11.5 min, 5% B; 12.5 min, 5% B. Data were acquired using DDA mode or full-scan mode in electrospray positive ionization mode.

For DDA mode, the parameters were set as: sheath gas flow 53 L/min, aux gas flow rate 14 L/min, sweep gas flow 3 L/min, spray voltage 3.5 kV, inlet capillary 269 °C, aux gas heater 438 °C, S-lens RF level 50.0. MS scan range was set as 100-1000 *m/z* with mass resolution of 35,000 at *m/z* 200. Automatic gain control (AGC) target was set to 1E6 with a maximum injection time of 100 ms. Up to 5 MS/MS spectra per MS1 were collected per cycle with mass resolution 17,500 at *m/z* 200, maximum injection time of 150 ms with an AGC target of 5E5. Isolation window was set to 1 *m/z* and the isolation offset at 0 *m/z*. Stepwised normalized collision energies were set at 25 eV, 40 eV, and 60 eV. The apex trigger was set to 2-15 s and a dynamic exclusion of 5 s. Isotopes were excluded from the analysis.

For full-scan mode, the parameters were set as: sheath gas flow 53 L/min, aux gas flow rate 14 L/min, sweep gas flow 3 L/min, spray voltage 3.5 kV, inlet capillary 269 °C, and aux gas heater 438 °C. MS scan range was set as 100-1000 *m/z* with mass resolution of 70,000 at *m/z* 200. AGC target was set to 1E6 with maximum injection time as 150 ms. Data in full-scan mode were acquired using different isCID energies: 0 eV, 10 eV, and 20 eV.

### Statistical analysis

For the IBD dataset, missing values were filled using the minimum of 5E5 and 10% of the minimum intensity for each feature. Outlier removal was conducted using the interquartile range (IQR) method. Data points below Q1 − 1.5 × IQR or above Q3 + 1.5 × IQR were removed from each group (non-IBD, CD or UC). Two-sided Mann-Whitney *U* tests were performed. For t-SNE visualization, intensity values were subjected to log transformation and feature-wise z-normalization.

### Molecular networking

A molecular network was constructed using the pseudo MS/MS spectra from the IBD dataset (HILIC positive mode). For the purpose of molecular networking, we cleaned each metabolic feature’s pseudo MS/MS by removing all ions larger than the feature *m/z* to enable modified cosine calculations. This cleaning step was not performed for the ms1-id annotation process, which utilizes the full pseudo MS/MS spectra with the open search approach described previously. Then, an MGF file for all cleaned pseudo MS/MS spectra was prepared. The mgf file was uploaded onto the GNPS2 platform, where a classical molecular networking workflow (version 2024.09.20) was completed. A minimum of modified cosine of 0.8 and matched peaks of 6 are required to build an edge in the network construction. The job is available at https://gnps2.org/status?task=670aa34a07544a5cbbd1f1d40605f50f.

## Data availability

All the source data used in this study are publicly accessible.

LC-MS data: pooled chemical standards (GNPS/MassIVE, MSV000095789); NIST human feces (GNPS/MassIVE, MSV000095787); IBD dataset (Metabolomics Workbench, PR000639); mouse feces (GNPS/MassIVE, MSV000095868); microbe isolates (GNPS/MassIVE, MSV000090053; GNPS/MassIVE, MSV000085024; GNPS/MassIVE, MSV000090030; GNPS/MassIVE, MSV000090000); environmental DOM (GNPS/MassIVE, MSV000094338; GNPS/MassIVE, MSV000083888; GNPS/MassIVE, MSV000083632); plant extracts (GNPS/MassIVE, MSV000090975; GNPS/MassIVE, MSV000090968).

MS imaging data: mouse liver with spotted chemical standards (METASPACE, dataset ID 2020-12-07_03h16m14s); mouse brain (MetaboLights, MTBLS313); mouse body (METASPACE, dataset ID 2022-07-08_20h45m00s); human hepatocytes (METASPACE, project ID Rappez_2021_SpaceM); HeLa and NIH3T3 cells (METASPACE, project ID Rappez_2021_SpaceM); plant root (METASPACE, dataset ID 2025-01-07_19h33m53s); human liver (METASPACE, dataset ID 2017-11-28_11h28m57s); human kidney (METASPACE, dataset ID 2024-09-19_00h01m48s); mouse kidney (METASPACE, dataset ID 2019-03-28_18h03m06s); mouse brain MALDI-TOF (METASPACE, dataset ID 2024-12-21_10h17m55s).

## Code availability

The **ms1-id** Python package is available on PyPI. Source codes can be found at GitHub (https://github.com/Philipbear/ms1_id) and Zenodo (https://zenodo.org/records/14873172) under the Apache-2.0 license.

## Acknowledgement

This work was supported by BBSRC/NSF award 2152526 and National Institute of Health Sciences U24DK133658. We acknowledge the NIST Complex Microbial Systems Group for providing the NIST material. V.C.L. is supported by Fonds de recherche du Québec - Santé (FRQS) Postdoctoral fellowship (335368). O.F. is funded by National Institute of Health R01 GM155383.

## Discloses

P.C.D. is a scientific advisor and holds equity in Sirenas, Cybele, and bileOmix, and is a Scientific Co-founder, and advisor and holds equity in Ometa, Arome, and Enveda with prior approval by UC-San Diego. T.A. has a patent application on single-cell metabolomics, leads creation of a startup on single-cell metabolomics incubated at the BioInnovation Institute (BII) in Copenhagen, Denmark, and has a consultancy contract with BII.

## Author contributions

P.C.D. and S.X. conceived the research project. S.X. developed the computational algorithm and performed data analysis. V.C.L. collected the LC-MS data. M.E. collected the MS imaging data. Y.E. provided LC-MS file summaries in public repositories. H.Y. and O.F. developed MassCube. T.A. helped with MS imaging data analysis and validation. S.X. and P.C.D. drafted the manuscript. P.C.D. supervised the project. All authors approved the manuscript.

## References

1. Sud, M. et al. Metabolomics Workbench: An international repository for metabolomics data and metadata, metabolite standards, protocols, tutorials and training, and analysis tools. Nucleic Acids Res. 44, D463–D470 (2016).

2. Yurekten, O. et al. MetaboLights: open data repository for metabolomics. Nucleic Acids Res. 52, D640–D646 (2024).

3. Wang, M. et al. Sharing and community curation of mass spectrometry data with Global Natural Products Social Molecular Networking. Nat. Biotechnol. 34, 828–837 (2016).

4. Palmer, A. et al. FDR-controlled metabolite annotation for high-resolution imaging mass spectrometry. Nat. Methods 14, 57–60 (2017).

5. da Silva, R. R., Dorrestein, P. C. & Quinn, R. A. Illuminating the dark matter in metabolomics. Proc. Natl. Acad. Sci. 112, 12549–12550 (2015).

6. Blaženović, I. et al. Structure Annotation of All Mass Spectra in Untargeted Metabolomics. Anal. Chem. 91, 2155–2162 (2019).

7. Aksenov, A. A., da Silva, R., Knight, R., Lopes, N. P. & Dorrestein, P. C. Global chemical analysis of biology by mass spectrometry. Nat. Rev. Chem. 1, 1–20 (2017).

8. Bittremieux, W., Wang, M. & Dorrestein, P. C. The critical role that spectral libraries play in capturing the metabolomics community knowledge. Metabolomics 18, 94 (2022).

9. Abiead, Y. E. et al. Enabling pan-repository reanalysis for big data science of public metabolomics data. Preprint at 10.26434/chemrxiv-2024-jt46s (2024).

10. Dührkop, K. et al. SIRIUS 4: a rapid tool for turning tandem mass spectra into metabolite structure information. Nat. Methods 16, 299–302 (2019).

11. Xing, S., Shen, S., Xu, B., Li, X. & Huan, T. BUDDY: molecular formula discovery via bottom-up MS/MS interrogation. Nat. Methods 1–10 (2023) doi:10.1038/s41592-023-01850-x.

12. Gabelica, V. & Pauw, E. D. Internal energy and fragmentation of ions produced in electrospray sources. Mass Spectrom. Rev. 24, 566–587 (2005).

13. Abrankó, L., García-Reyes, J. F. & Molina-Díaz, A. In-source fragmentation and accurate mass analysis of multiclass flavonoid conjugates by electrospray ionization time-of-flight mass spectrometry. J. Mass Spectrom. 46, 478–488 (2011).

14. Criscuolo, A., Zeller, M. & Fedorova, M. Evaluation of Lipid In-Source Fragmentation on Different Orbitrap-based Mass Spectrometers. J. Am. Soc. Mass Spectrom. 31, 463–466 (2020).

15. Xu, Y.-F., Lu, W. & Rabinowitz, J. D. Avoiding Misannotation of In-Source Fragmentation Products as Cellular Metabolites in Liquid Chromatography–Mass Spectrometry-Based Metabolomics. Anal. Chem. 87, 2273–2281 (2015).

16. Giera, M., Aisporna, A., Uritboonthai, W. & Siuzdak, G. The hidden impact of in-source fragmentation in metabolic and chemical mass spectrometry data interpretation. Nat. Metab. 1–2 (2024) doi:10.1038/s42255-024-01076-x.

17. Domingo-Almenara, X. et al. Autonomous METLIN-Guided In-source Fragment Annotation for Untargeted Metabolomics. Anal. Chem. 91, 3246–3253 (2019).

18. Baygi, S. F., Kumar, Y. & Barupal, D. K. IDSL.CSA: Composite Spectra Analysis for Chemical Annotation of Untargeted Metabolomics Datasets. Anal. Chem. 95, 9480–9487 (2023).

19. Xue, J. et al. EISA-EXPOSOME: One Highly Sensitive and Autonomous Exposomic Platform with Enhanced in-Source Fragmentation/Annotation. Anal. Chem. 95, 17228–17237 (2023).

20. Wang, X.-C. et al. AntDAS-DDA: A New Platform for Data-Dependent Acquisition Mode-Based Untargeted Metabolomic Profiling Analysis with Advantage of Recognizing Insource Fragment Ions to Improve Compound Identification. Anal. Chem. 95, 638–649 (2023).

21. Wasito, H., Causon, T. & Hann, S. Alternating in-source fragmentation with single-stage high-resolution mass spectrometry with high annotation confidence in non-targeted metabolomics. Talanta 236, 122828 (2022).

22. Broeckling, C. D., Afsar, F. A., Neumann, S., Ben-Hur, A. & Prenni, J. E. RAMClust: A Novel Feature Clustering Method Enables Spectral-Matching-Based Annotation for Metabolomics Data. Anal. Chem. 86, 6812–6817 (2014).

23. Graça, G. et al. Automated Annotation of Untargeted All-Ion Fragmentation LC–MS Metabolomics Data with MetaboAnnotatoR. Anal. Chem. 94, 3446–3455 (2022).

24. Kachman, M. et al. Deep annotation of untargeted LC-MS metabolomics data with Binner. Bioinformatics 36, 1801–1806 (2020).

25. Li, Y. et al. Spectral entropy outperforms MS/MS dot product similarity for small-molecule compound identification. Nat. Methods 18, 1524–1531 (2021).

26. Aksenov, A. A. et al. Auto-deconvolution and molecular networking of gas chromatography–mass spectrometry data. Nat. Biotechnol. 39, 169–173 (2021).

27. Xue, J. et al. Enhanced in-Source Fragmentation Annotation Enables Novel Data Independent Acquisition and Autonomous METLIN Molecular Identification. Anal. Chem. 92, 6051–6059 (2020).

28. Wishart, D. S. et al. HMDB 5.0: the Human Metabolome Database for 2022. Nucleic Acids Res. 50, D622–D631 (2022).

29. Sud, M. et al. LMSD: LIPID MAPS structure database. Nucleic Acids Res. 35, D527– D532 (2007).

30. Hastings, J. et al. ChEBI in 2016: Improved services and an expanding collection of metabolites. Nucleic Acids Res. 44, D1214–D1219 (2016).

31. Schymanski, E. L. et al. Identifying Small Molecules via High Resolution Mass Spectrometry: Communicating Confidence. Environ. Sci. Technol. 48, 2097–2098 (2014).

32. Tsugawa, H. et al. MS-DIAL: data-independent MS/MS deconvolution for comprehensive metabolome analysis. Nat. Methods 12, 523–526 (2015).

33. Kuhl, C., Tautenhahn, R., Böttcher, C., Larson, T. R. & Neumann, S. CAMERA: An Integrated Strategy for Compound Spectra Extraction and Annotation of Liquid Chromatography/Mass Spectrometry Data Sets. Anal. Chem. 84, 283–289 (2012).

34. Mahieu, N. G. & Patti, G. J. Systems-Level Annotation of a Metabolomics Data Set Reduces 25 000 Features to Fewer than 1000 Unique Metabolites. Anal. Chem. 89, 10397–10406 (2017).

35. Nash, W. J., Ngere, J. B., Najdekr, L. & Dunn, W. B. Characterization of Electrospray Ionization Complexity in Untargeted Metabolomic Studies. Anal. Chem. 96, 10935–10942 (2024).

36. Abramson, F. P. Automated identification of mass spectra by the reverse search. Anal. Chem. 47, 45–49 (1975).

37. Djoumbou Feunang, Y., et al. ClassyFire: automated chemical classification with a comprehensive, computable taxonomy. J. Cheminformatics 8, 61 (2016).

38. Lloyd-Price, J. et al. Multi-omics of the gut microbial ecosystem in inflammatory bowel diseases. Nature 569, 655–662 (2019).

39. Kim, H. W. et al. NPClassifier: A Deep Neural Network-Based Structural Classification Tool for Natural Products. J. Nat. Prod. 84, 2795–2807 (2021).

40. Lavelle, A. & Sokol, H. Gut microbiota-derived metabolites as key actors in inflammatory bowel disease. Nat. Rev. Gastroenterol. Hepatol. 17, 223–237 (2020).

41. Chang, F.-Y. et al. Gut-inhabiting Clostridia build human GPCR ligands by conjugating neurotransmitters with diet- and human-derived fatty acids. Nat. Microbiol. 6, 792–805 (2021).

42. Cohen, L. J. et al. Commensal bacteria make GPCR ligands that mimic human signalling molecules. Nature 549, 48–53 (2017).

43. Derave, W., Everaert, I., Beeckman, S. & Baguet, A. Muscle Carnosine Metabolism and β-Alanine Supplementation in Relation to Exercise and Training. Sports Med. 40, 247–263 (2010).

44. Boldyrev, A. A., Aldini, G. & Derave, W. Physiology and Pathophysiology of Carnosine. Physiol. Rev. 93, 1803–1845 (2013).

45. Guney, Y. et al. Carnosine may reduce lung injury caused by radiation therapy. Med. Hypotheses 66, 957–959 (2006).

46. Rappez, L. et al. SpaceM reveals metabolic states of single cells. Nat. Methods 18, 799– 805 (2021).

47. Giné, R. et al. HERMES: a molecular-formula-oriented method to target the metabolome. Nat. Methods 18, 1370–1376 (2021).

48. Baquer, G. et al. rMSIfragment: improving MALDI-MSI lipidomics through automated in-source fragment annotation. J. Cheminformatics 15, 80 (2023).

49. Watrous, J., et al. Mass spectral molecular networking of living microbial colonies. Proc. Natl. Acad. Sci. 109, E1743–E1752 (2012).

50. Fiehn, O., et al. MassCube: a Python framework for end-to-end metabolomics data processing from raw files to phenotype classifiers. Preprint at 10.21203/rs.3.rs-5530740/v1 (2025).

51. Senan, O. et al. CliqueMS: a computational tool for annotating in-source metabolite ions from LC-MS untargeted metabolomics data based on a coelution similarity network. Bioinformatics 35, 4089–4097 (2019).

52. Uppal, K., Walker, D. I. & Jones, D. P. xMSannotator: An R Package for Network-Based Annotation of High-Resolution Metabolomics Data. Anal. Chem. 89, 1063–1067 (2017).

53. Wang, M. et al. Mass spectrometry searches using MASST. Nat. Biotechnol. 38, 23–26 (2020).

54. Zuffa, S. et al. microbeMASST: a taxonomically informed mass spectrometry search tool for microbial metabolomics data. Nat. Microbiol. 1–10 (2024) doi:10.1038/s41564-023-01575-9.

55. Gentry, E. C. et al. Reverse metabolomics for the discovery of chemical structures from humans. Nature 1–8 (2023) doi:10.1038/s41586-023-06906-8.

56. Bittremieux, W. et al. Open access repository-scale propagated nearest neighbor suspect spectral library for untargeted metabolomics. Nat. Commun. 14, 8488 (2023).

57. Lam, S. K., Pitrou, A. & Seibert, S. Numba: a LLVM-based Python JIT compiler. in Proceedings of the Second Workshop on the LLVM Compiler Infrastructure in HPC 1–6 (Association for Computing Machinery, New York, NY, USA, 2015). doi:10.1145/2833157.2833162.

58. Li, Y. & Fiehn, O. Flash entropy search to query all mass spectral libraries in real time. Nat. Methods 1–4 (2023) doi:10.1038/s41592-023-02012-9.

59. Coman, C. et al. Simultaneous Metabolite, Protein, Lipid Extraction (SIMPLEX): A Combinatorial Multimolecular Omics Approach for Systems Biology*. Mol. Cell. Proteomics 15, 1435–1466 (2016).

