## Supplementary Table 1 for "Structural annotation of full-scan MS data: A unified solution for LC-MS and MS imaging analyses"

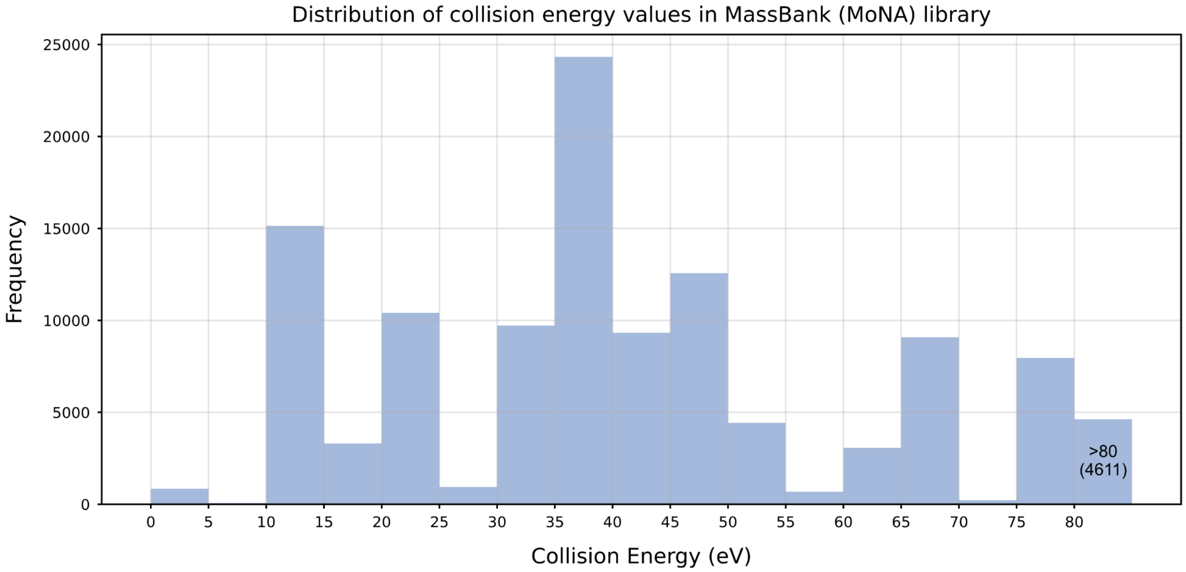


**Supplementary Fig. 1 |** Most reference MS/MS spectra are collected under medium to high collision energies (CE). Analysis of the public MoNA library reveals that reference spectra are predominantly acquired at CEs between 10-55 eV, with notable peaks at 15 eV and 35 eV. The distribution shows that only 0.81% of all reference spectra were collected under low collision energies (≤5 eV), highlighting a significant gap in spectral databases for low-energy fragmentation patterns.

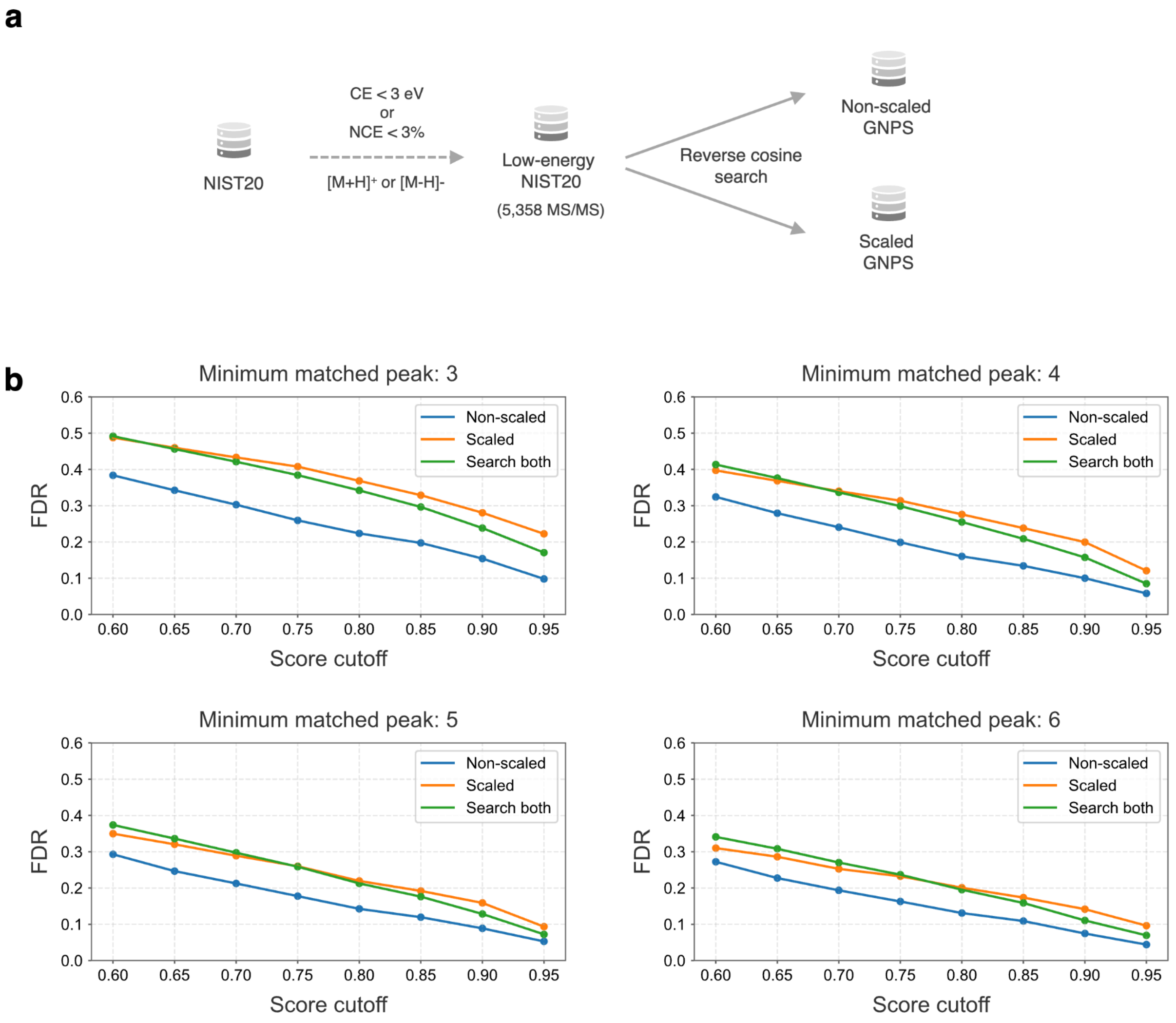


**Supplementary Fig. 2 |** Peak scaling tends to increase the false discovery rate (FDR) during spectral matching. We selected low-energy reference MS/MS spectra from NIST20 using collision energy (CE) < 3 eV or normalized collision energy (NCE) < 3%. These spectra were then searched against the non-scaled and scaled GNPS library. FDR-score plots with different minimum matched peaks were shown. We expect future optimization of scaling can further improve the annotation confidence.

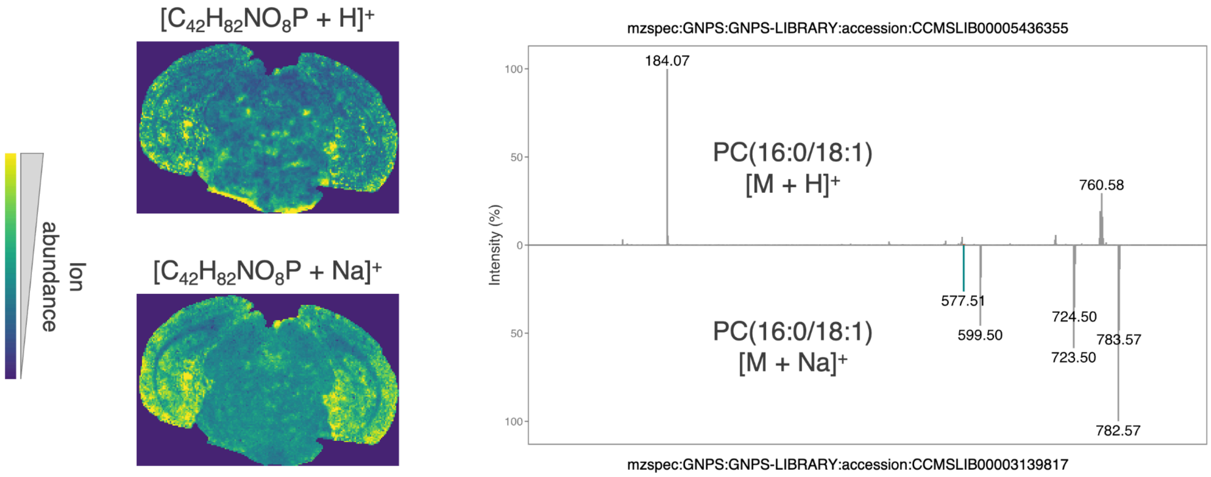


**Supplementary Fig. 3 |** PC(16:0/18:1) annotated in the mouse brain MS imaging data, in both $[M+H]^+$ and $[M+Na]^+$ ion forms. Distinct localization and fragmentation behaviors were shown.

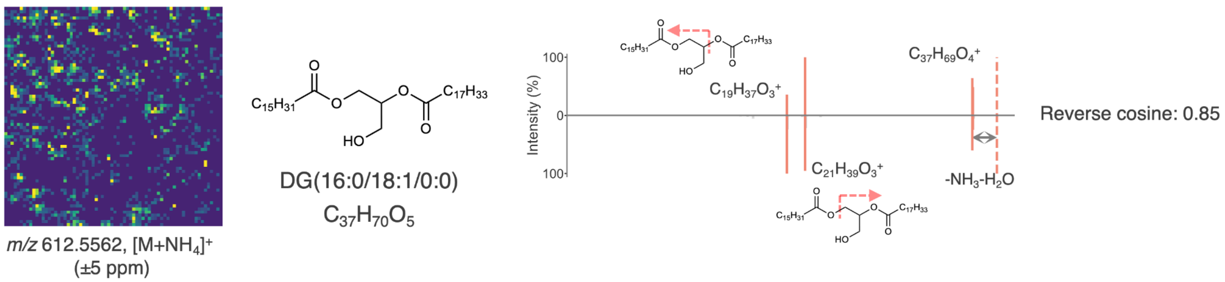


**Supplementary Fig. 4 |** DG(16:0/18:1/0:0) was annotated in hepatocytes MS imaging data, with a reverse cosine score of 0.85. Both acyl chains could be annotated in the MS/MS spectrum. The ion image was created using a mass tolerance of 5 ppm.

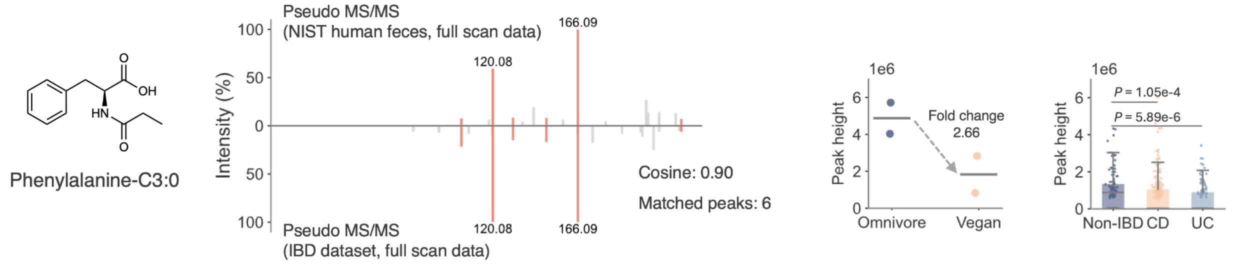


**Supplementary Fig. 5 |** A proof-of-principle of MS1-based MASST. We searched the pseudo MS/MS spectrum of phenylalanine-C3:0 from NIST human feces data against the pseudo MS/MS spectra pool from the IBD dataset, and it returned a match of cosine 0.90 with 6 matched peaks. The returned hit was also annotated as phenylalanine-C3:0 in the IBD dataset.

**Supplementary Table 1 |** LC-MS data file counts in three main metabolomics repositories.

| Repository name | Total file count | File count of MS1-only data | Percentage of MS1-only data |
| --- | --- | --- | --- |
| GNPS/MassIVE | 291080 | 43718 | 15.02% |
| MetaboLights | 115138 | 99600 | 86.50% |
| Metabolomics Workbench | 58458 | 46836 | 80.12% |
| **Total** | **464676** | **190154** | **40.92%** |
